# Enabling Real-time Process Analysis in Embedded Bioprinting with a Modular In Situ Monitoring Platform

**DOI:** 10.1101/2025.06.25.661611

**Authors:** Giovanni Zanderigo, Ferdows Afghah, Bianca Maria Colosimo, Ritu Raman

## Abstract

Real-time monitoring and *in situ* data analysis are increasingly vital for enhancing precision, reproducibility, and defect detection in embedded bioprinting. As interest grows in improving the capabilities of existing bioprinting systems, accessible strategies for integrating real-time sensing and analysis are becoming essential to ensure consistent quality and to optimize printing parameters. Here, we present a modular, low-cost, and printer-agnostic platform that combines a compact sensing architecture with an effective image analysis pipeline to enable *in situ* process monitoring, defect detection, and print quality assessment. the platform integrates a digital microscope aligned on-axis with the extrusion printhead to capture high-resolution images during fabrication. We applied and compared two segmentation methods, thresholding and the Segment Anything Model (SAM) on *in situ* and *ex situ* datasets acquired via confocal fluorescence imaging, finding SAM to yield stronger correlations (R = 0.85–0.86) between *in situ* and *ex situ* measurements. Additionally, we demonstrated that 2D *in situ* images provide reliable approximations of 3D filament geometries, supporting their use for real-time morphological assessment. the system also revealed pressure-related effects on the diameters, and a critical velocity threshold for printing stability, highlighting its value for process optimization. together, these findings establish the approach as a low-cost, scalable and adaptable solution that can be readily implemented across embedded bioprinting workflows, offering a practical path toward greater reproducibility and automation.

## Introduction

Three-dimensional (3D) bioprinting enables the fabrication of multicellular constructs using various biomaterials, cells, and biomolecules ^1–3^. Embedded bioprinting, which involves depositing bioinks within a viscoelastic support material, expands extrusion capabilities ^4,5^. the elastic properties of the support material allow smooth needle motion, and deposition of soft bioinks, while the viscous behavior retains the deposited bioink, enabling the fabrication of complex and overhanging structures. Despite these advantages, extrusion-based bioprinting remain prone to printing defects such as over-extrusion, which leads to excessive material deposition and structural loss, and under-extrusion, which results in filament discontinuities and weak mechanical integrity. These errors compromise the structural and functional fidelity of engineered tissues, particularly where fine architectural features are critical ^6,7^. Real-time monitoring can help detect such defects early and enable dynamic adjustments of printing parameters such as speed and pressure to stabilize the process. However, there is a notable lack of affordable, modular solutions for *in situ* monitoring that can be easily integrated into standard bioprinting platforms ^8–10^.

Real-time control over bioprinting process parameters requires methods for *in situ* monitoring of 3D bioprinting. By addressing potential flaws and inaccuracies in the final structure, *in situ* monitoring could enhance the consistency and reproducibility of the printing process, as well as increase the structural integrity of printed tissues ^11–13^. Enabling quality control during printing could also drastically reduce time and material waste as compared to iterative experimental optimization.

Over the past decade, several approaches have been proposed in the literature for *in situ* monitoring of conventional additive manufacturing processes involving metals and polymers ^14–16^. More recently, attention has shifted toward adapting these solutions for bioprinting as well^7,17–19^, with the aim of assessing shape fidelity by detecting irregularities, process instabilities, or local defects along the tool path. Among the sensing architectures explored in this context, the most commonly investigated include optical imaging^18,20^, thermal imaging using infrared (IR) cameras ^7,21,22^, optical coherence tomography (OCt) ^7,12,23^, fluorescence imaging ^24^, and laser scanning ^25,26^. Each monitoring technique presents its own advantages and limitations. For instance, while OCt offers high spatial resolution, it suffers from limited depth penetration, requires complex data processing, and entails high equipment costs ^14,25^. Conversely, the spatial and temporal resolution of conventional cameras may be insufficient—unless high-end, costly solutions are used.

Moreover, the transparency of many bioinks can hinder visible light imaging, making thermal cameras operating in the infrared (IR) range a more suitable alternative ^7^.

With reference to embedded bioprinting, *in situ* monitoring remained largely unexplored until a few recent attempts. In one such study, researchers employed OCT for embedded bioprinting by developing a custom setup ^27^. To address the limitations of OCT, the support bath formulation was modified in order to increase its transparency, and printed it sequentially with the biomaterial, adding complexity to the fabrication process. In another study, researchers integrated a high-resolution camera with a custom 3D bioprinting platform to monitor the embedded extrusion process ^28^. Their setup employed an additional motorized system that positioned the camera beneath the print bead, using a glass platform to allow imaging from below. However, this configuration requires a custom bioprinter and limits accessibility for imaging multi-layered constructs. The fact that these approaches often demand complex bath reformulations, hardware modifications, or advanced software integration, highlights the need for a modular, cost-effective system that is easy to retrofit into existing bioprinters and does not require invasive changes to the materials or printing setup.

Here, we introduce a visible-light-based *in situ* monitoring platform built around a commercially available digital microscope aligned with the printhead. The system captures high-resolution images during embedded bioprinting without requiring modifications to the support bath or the use of additional dyes. We validate our approach against confocal fluorescence microscopy (Figure 1a–b), comparing two segmentation strategies: traditional thresholding for edge detection and a recent AI-based foundation model—the Segment Anything Model (SAM), a promptable vision model designed for general-purpose object segmentation. Our proposed solution is capable of revealing critical process thresholds and systematic geometric deviations, making it a promising tool for real-time process understanding and the development of future closed-loop control strategies.

**Figure 1.**
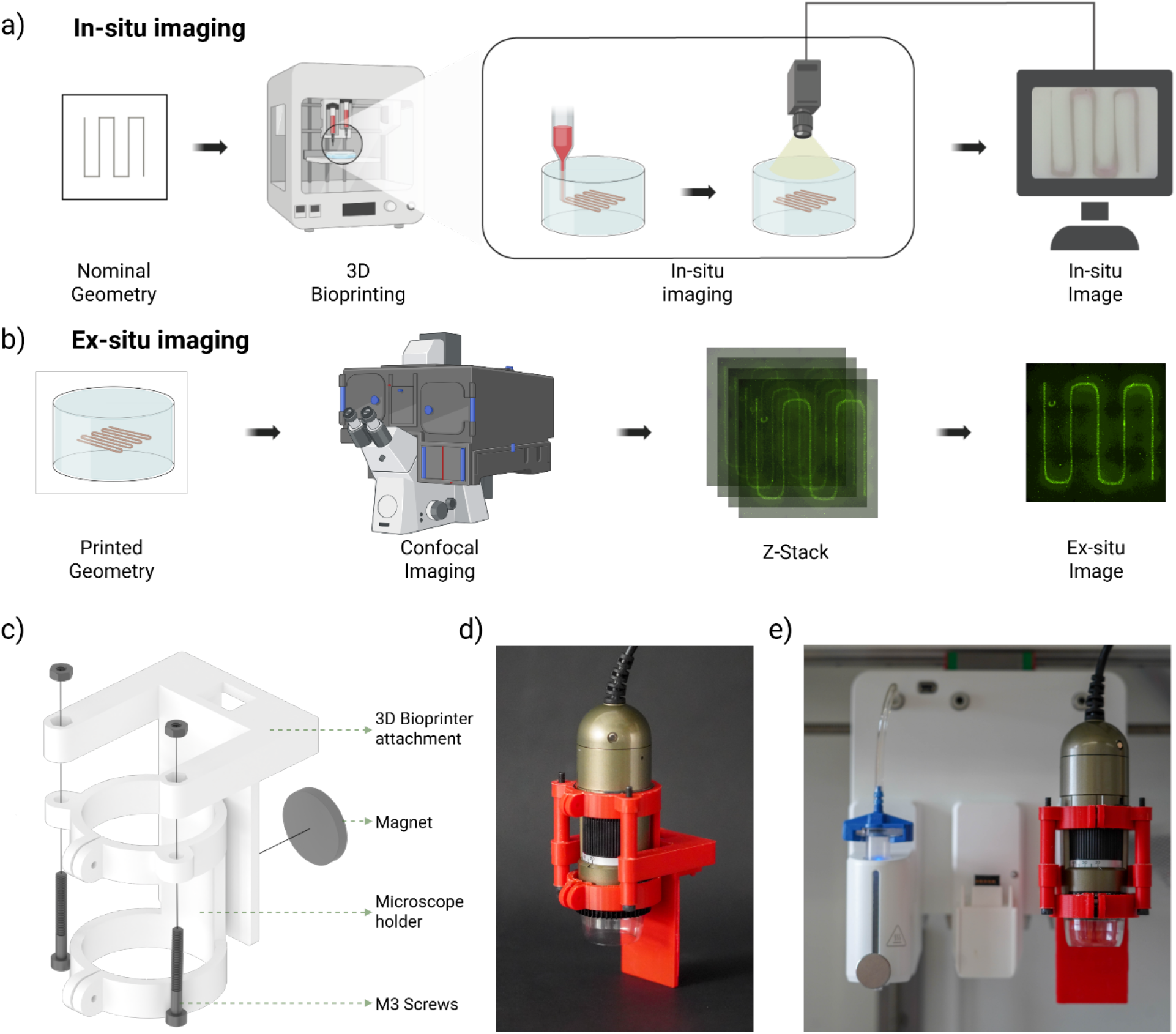
**a)** Schematic of the *in situ* imaging process, which involves printing the desired geometry layer by layer; after each layer is completed, the printhead moves aside, allowing the microscope to position itself directly above the printed structure and capture a high-resolution image; **b)** Schematic of the *ex situ* imaging workflow using confocal microscopy, in which the same constructs imaged *in situ* are subsequently analyzed with fluorescence confocal microscopy to obtain high-resolution z-stacks; stacks were then collapsed in a max projection image used as final *ex situ* image; **c)** Computer-aided design (CAD) model of the custom digital microscope support; **d)** View of the Dino-Lite digital microscope housed within the custom 3D printed support; **e)** Image of digital microscope assembly mounted on the bioprinter printhead together with the extruder.

## Results

### Modular in situ monitoring system

To enable reliable real-time visualization of embedded bioprinting, we developed a compact, modular *in situ* imaging system that captures high-resolution images during fabrication, allowing immediate defect detection and print quality assessment.

The system integrates a Dino-Lite digital microscope onto the printhead assembly, offering a resolution of 1.3 megapixels, with a magnification range of 10x-90x. The custom holder for the microscope was designed for ease of assembly using low-cost components and fabricated via extrusion-based 3D printing (Figure 1c-d). It attaches securely to an existing printhead slot on a standard commercial multi-material extrusion bioprinter through an interlocking mechanism reinforced with an embedded magnet, requiring only standard M3 screws for assembly (Figure 1e). By leveraging the controlled xyz motion of the printhead, the system maintains a fixed working distance, enabling consistent automatic image acquisition after each printed layer. A USB-powered LED panel (MSAK827, DUNWELL TECH) was positioned beneath the printing substrate (Petri dish) to enhance contrast and visibility.

The sensor was aligned along the axis perpendicular to the bath surface to effectively addresses the issue of surface reflections that often arise with suspension baths, minimizing glare and enhancing image clarity during imaging. Given this spatial constraint, the system addresses the confined space within bioprinters and the need to avoid interference with moving components with a highly compact and non-intrusive design. The modularity of the system allows it to be mounted and adjusted around existing hardware with minimal modification, ensuring compatibility with a variety of printer architectures and digital microscopes.

### Experimental Setup and Integration of Monitoring System

To systematically investigate how printing speed affects construct fidelity and process stability for embedded bioprinting, we conducted a controlled experiment where all variables were kept constant except for print velocity. Speed was selected as the key variable due to its critical role in determining print fidelity and the amount of material deposited, directly influencing under and over extrusion.

Constructs were printed at 6 different printing velocities, namely 3, 5, 7, 9, 11, and 13 mm/s, each tested in triplicate and printed in randomized order ^29^. Pneumatic pressure was kept constant at 15 kPa, based on preliminary trials that identified it as optimal for extrusion. A 10×10×0.4 mm coil-like geometry was printed (Figure 1a), incorporating both long (vertical) and short (horizontal) filaments with curved transitions. This design enabled evaluation of printing performance under both stationary conditions characterized by long rectilinear filaments, and dynamic ones characterized by short filaments and rapid velocity transitions.

The Support bath formulation used for embedded printing consisted of commonly used components, Pluronic F127, laponite, and calcium chloride with no modifications such as added dyes or transparency enhances, preserving the standard bioprinting conditions typically reported in the literature ^30^. The bioink was based on fibrin, which exhibited intrinsic auto fluoresces, enabling imaging without requiring additional fluorescent labeling or compromising the mechanical and rheological properties of the material. *In situ* images were captured during each print using the integrated monitoring system, allowing real-time visualization of the process (Figure 1a). After printing, all the structures were then imaged *ex situ* with a confocal fluorescence microscope (Figure 1b). This resulted in matching *in situ* and *ex situ* image pairs for each construct, captured during and after printing, respectively. The high-resolution *ex situ* images served as a reference to validate the measurements and structural information extracted from *in situ* data.

### Integrating Enhanced Image Analysis for Process Modeling and Monitoring

To enable quantitative comparison of the printed constructs, image segmentation was performed on both *in situ* and *ex situ* images to extract relevant geometric features. Binary segmentation masks were generated using both a classic approach based on thresholding of the grey-scale intensities, and the recent AI-based foundation model Segment Anything Model (SAM) solution developed by Meta AI ^31^ (Figure 2).

**Figure 2.**
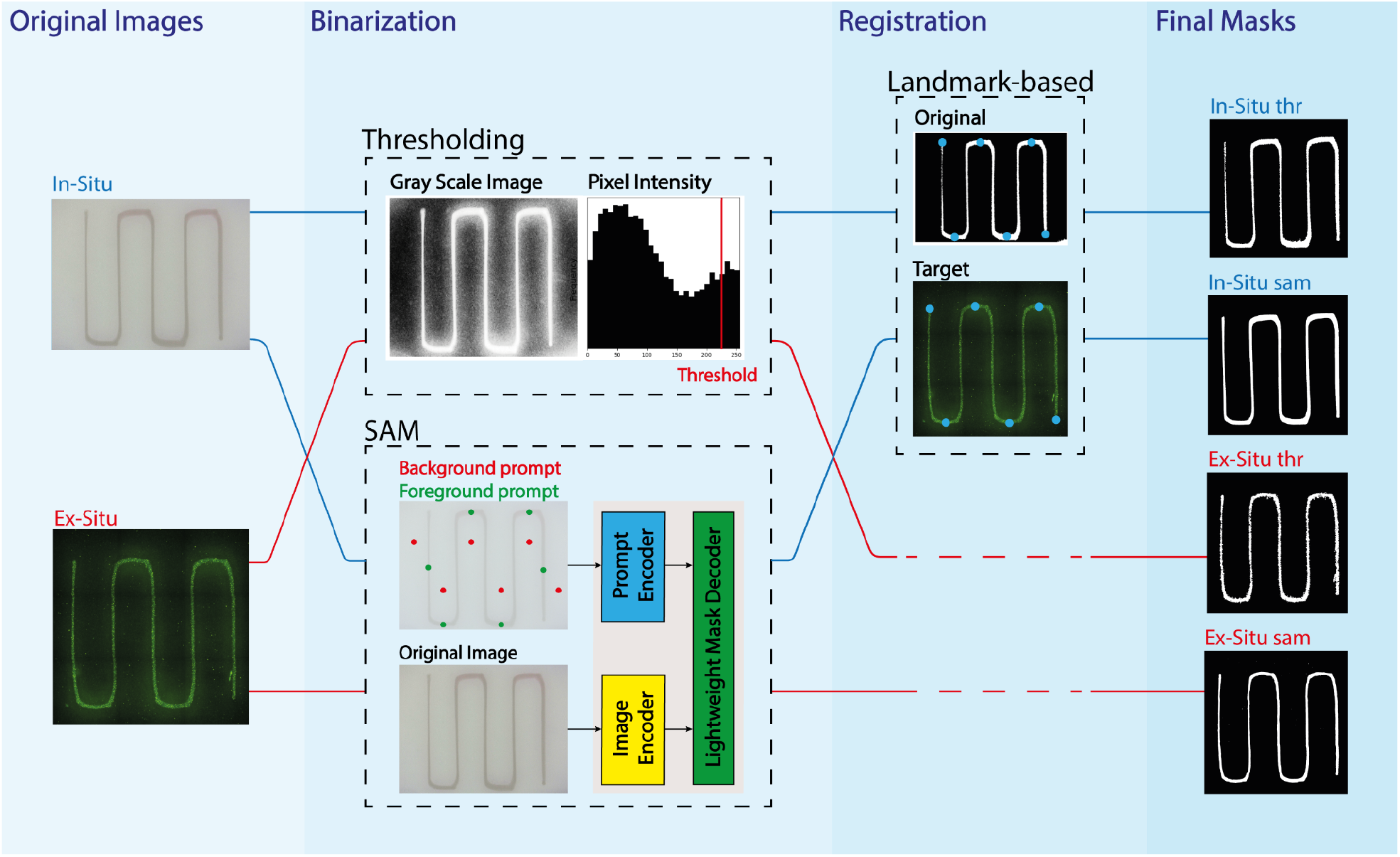
Image segmentation process schematic. From left to right: examples of *in situ* and *ex situ* original images used for the binarization; binarization pipelines adopted on both the *in situ* and *ex situ* images using thresholding (top) and SEM (bottom); registration of the *in situ* images over the *ex situ* images to achieve the same resolution (5940×5940) and making them suitable for overlay analysis adopting a landmark-based registration algorithm; example of the two types of final masks obtained with the two segmentation algorithms, both for *in situ* and *ex situ* (for a total of 4 different masks).

Although conceptually different, both approaches aim to generate binary masks distinguishing foreground from background pixels. The thresholding method starts with grayscale conversion of the image and exploits intensity differences between object and background. Based on pixel intensity histograms, various thresholding methods exist ^32,33^. In our case, a fixed threshold (pixel intensity of 235) was used as it outperformed competitor solutions and was therefore selected to ensure consistent preprocessing and reproducible segmentation across all 18 images (Figure 2).

In contrast, SAM is a vision transformer model pre-trained on large datasets ^31^, enabling few-shot learning for segmentation due to its prompt-based architecture, that has already demonstrated high potential in scientific application ^34^. Prompts were generated by selecting foreground and background points, then passed to the model along with the image to output a binary mask (Figure 2). For this purpose, a specific pipeline was developed on Python, able to collect input images, and then automatically generate prompts by selecting positive and negative points over the images to define foreground and background regions. Images and prompt were then passed to the last checkpoint of the model (sam_vit_h_4b8939.pth) that was downloaded from GitHub. Foreground prompts were placed along the G-code-defined geometry, while background points were set between filaments. This setup avoids manual annotation while still guiding SAM effectively, making it a practical zero-shot solution for *in situ* monitoring. The prompt strategy was uniformly applied to all images. This method can be easily implemented in 3D printing workflows thanks to the G-code-based nature of the process, and although its computational time is higher than simple thresholding techniques, it remains reasonable within the overall timeline of a live monitoring system for embedded bioprinting.

Finally, all *in situ* masks were registered to their corresponding *ex situ* images (5940×5940) to enable resolution matching and overlay analysis.

### Validation of 2D Geometry Against 3D Structure

To validate whether 2D image projection can readily represent 3D filament structures, 3D stacks acquired via confocal microscopy were analyzed using the software IMARIS. The analysis consisted in the initial reconstruction of the filament surface from the fibrin fluorescence signal (Figure 3a). Three cross sections were then taken from the reconstructed volume, and the circularity of the filaments was assessed by computing the ratio of the height over the width of cross-sectional profiles along the filament length. Results confirmed minimal distortion in projection, since the average ratio was (*H*/*W*)_*avg*_ = 1.003 ± 0.117 over the three cross sections of the filament, supporting the use of 2D *in situ* images for reliable 3D geometric quantification.

**Figure 3.**
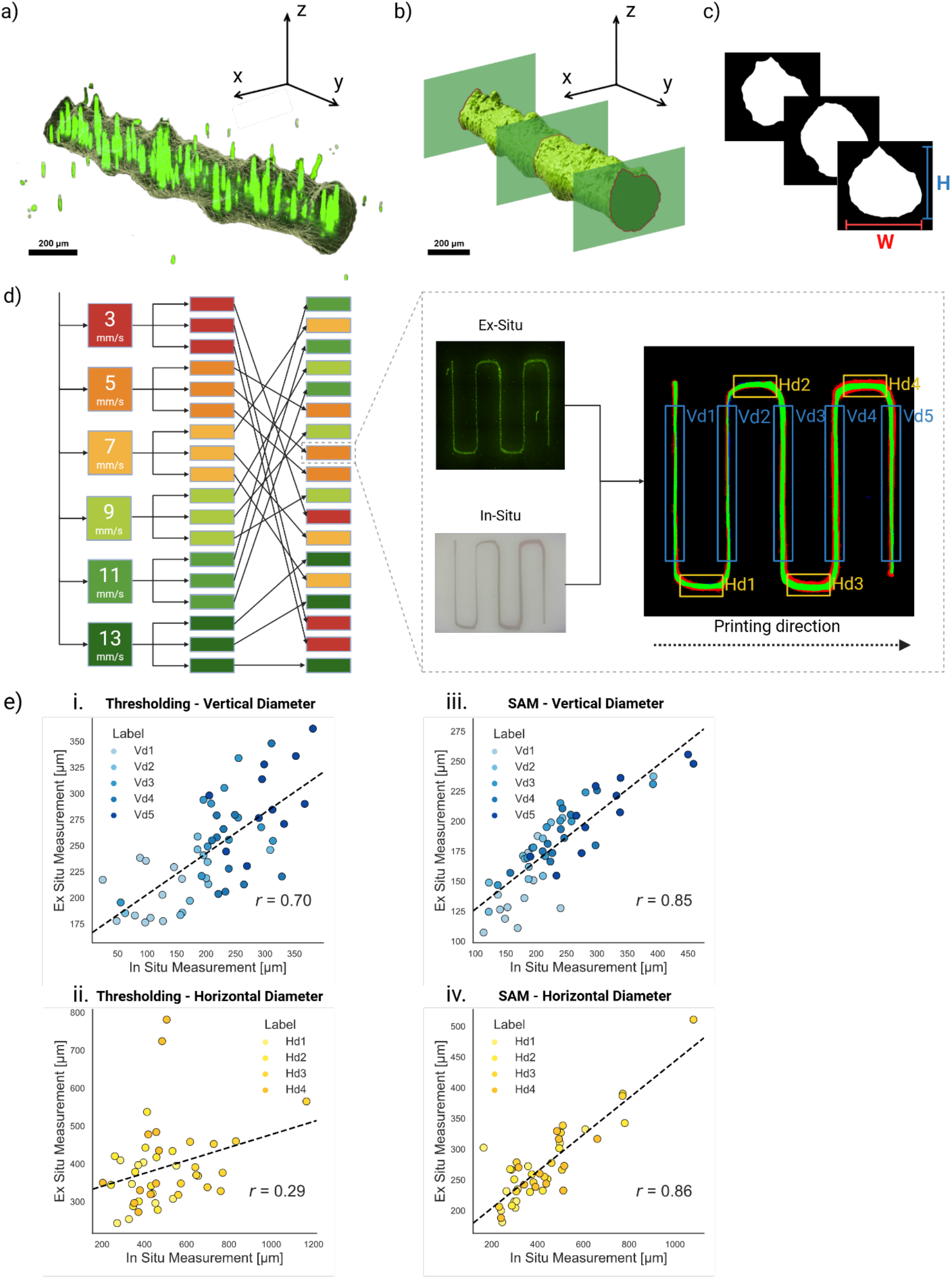
**a)** Surface reconstruction performed on IMARIS starting from the bio-ink fluorescent signal collected with confocal imaging; **b)** Extraction of the three different cross-sectional planes along the filament length; **c)** Estimation of height-to-width (H/W) ratio of the sections to assess circularity in 2D projections; **d)** Schematic showing the range of printing velocities used in this study; the triplicated experiments; the randomization to get rid of nuisance factors that could bias the result; the extracted overlay images used to estimate the *in situ* and the *ex situ* diameters values; **e)** Correlation analysis of the *in situ* and *ex situ* measurements made with: **i)** the thresholding on vertical diameters, **ii)** thresholding on horizontal diameters, **iii)** SAM on vertical diameters, **iv)** SAM on horizontal diameters.

### Validation of the Segmentation Approaches through In situ and Ex situ Diameter Correlation

A correlation analysis was used to assess how well *in situ* image-based filament diameters matched *ex situ* ground truth data, facilitating the validation of segmentation methods. From each image, 5 vertical and 4 horizontal Regions Of Interest (ROIs) were extracted to estimate the vertical and horizontal diameters. Each ROI was aligned vertically by performing a Principal Component Analysis ^35^ on the pixels of the mask, and the diameters of the ROIs were extracted. The diameters were ordered based on the printing direction from Vd1 to Vd5 for the vertical sections, and from Hd1 to Hd4 for the horizontal (Figure 3d).

In the correlation analysis, both segmentation approaches were compared to assess their suitability for this task. Initially, all data points were included, but outliers from defect-prone points referred to as “spilled prints” due to visible material overflow into the support bath-lowered the correlation for both segmentation approaches. These spills typically occurred at lower printing velocities, where excess material failed to form stable filaments and instead accumulated in irregular shapes, as further examined in the following sections. After removing these defect-related data points, cleaned datasets were used for analysis (Figure 3e).

A correlation of R = 0.70 was observed for the vertical diameters extracted from the thresholding masks (Figure 3e-i), while the horizontal segments showed a lower correlation of R = 0.29 (Figure 3e-ii). This discrepancy can be primarily attributed to reduced contrast in the curved regions of the filaments, where the thresholding segmentation lacks precision and tends to overestimate *in situ* diameters. For the diameters extracted with SAM, the correlations were notably higher, with R = 0.85 for the vertical (Figure 3e-iii) and R = 0.86 for the horizontal filaments (Figure 3e-iv). Although the *in situ* measurements consistently yielded slightly larger diameter estimates compared to the *ex situ* ones, the agreement between the two methods was much stronger, as reflected by the higher correlation coefficients. This overestimation averaged approximately 27% for vertical filaments and 53% for horizontal ones, potentially due to refraction or optical blurring caused by the bath interface. It should be noted, however, that these overestimation values are consistent, given the correlation of 0.85 and 0.86, and therefore simply represent a measurement bias. Mitigation strategies could include calibrating the system with the known overestimation or applying image deconvolution techniques to compensate for systematic optical distortions.

### Print Quality and Process Optimization Enabled by In Situ Monitoring Setup

Having established the reliability of the *in situ* SAM measurements, we used them to investigate how print velocity and extrusion dynamics can affect print quality and shape fidelity in embedded bioprinting.

We first analyzed the correlation between printing velocity and diameter of both vertical (Figure 4a-b) and horizontal (Figure 4c-d) filaments. Results showed a strong influence of velocity: at low speeds, failures due to spilling were frequent—100% at 3 mm/s (3 of 3 prints) and 67% at 5 and 7 mm/s (2 of 3). As previously anticipated, the spilling typically occurred at lower printing velocities, where the excessive amount of deposited material within the support bath was not retained within the gel structure, but rather spilled over the surface of the support bath. In contrast, from 9 mm/s upward, prints were free of spill and showed stable diameters, suggesting 9 mm/s is a critical threshold for stable, reproducible printing. This behavior is likely attributed to the size of the 0.33 mm needle used, which limits gel displacement and leads to the formation of filaments with consistent diameters.

**Figure 4.**
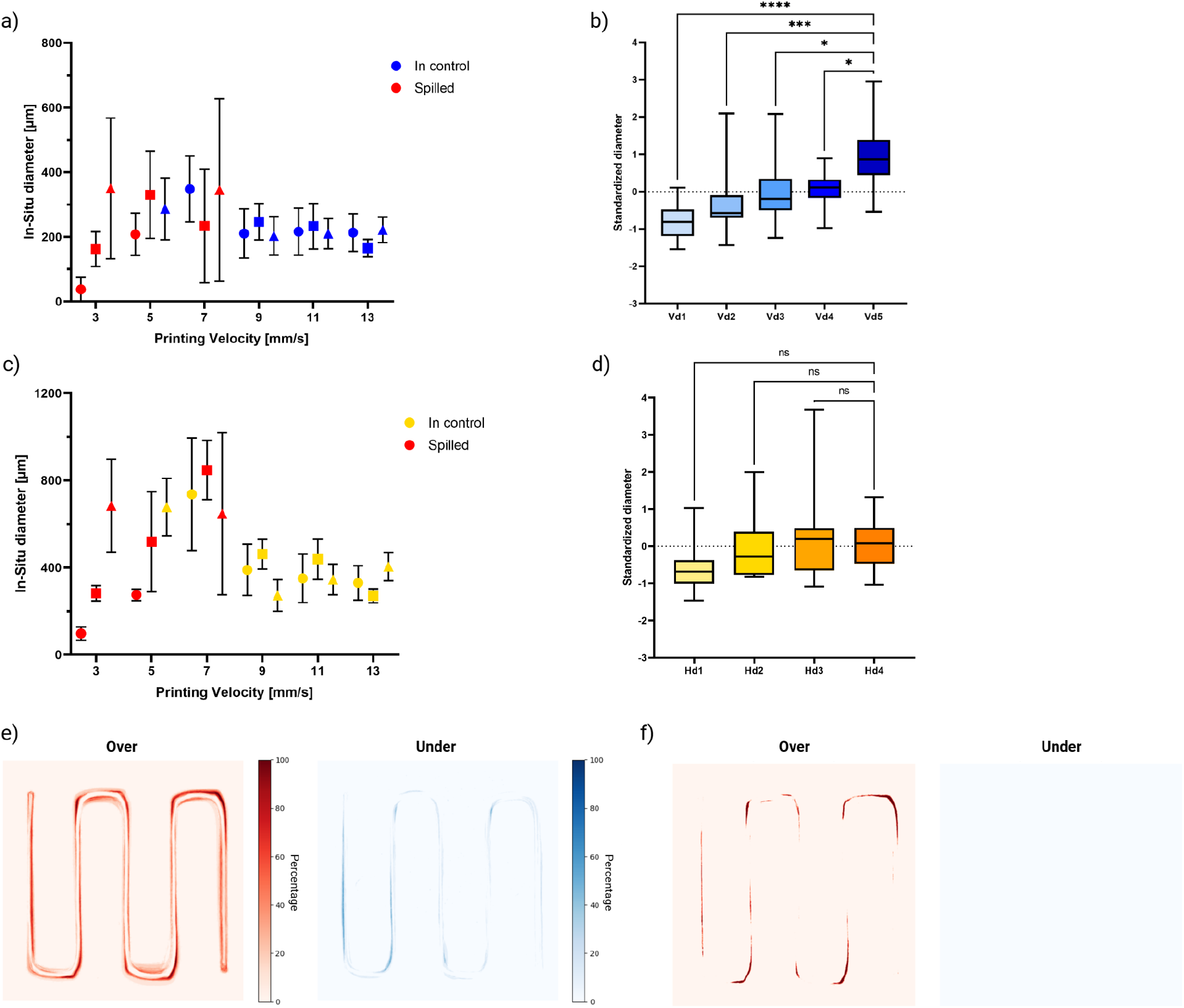
**a)** Scatterplot of the vertical filament diameter grouped by the printing velocity showing lager variance below 9mm/s and a stable performance above the same velocity; **b)** Box plot of the standardized vertical diameters grouped across sequential regions (Vd1-Vd5), showing the increasing trend in diameter within the same prints; **c)** Scatterplot of horizontal filament diameter as a function of printing velocity also showing lager variance below 9mm/s and a stable performance above the same velocity; **d)** Box plot of the standardized horizontal diameters grouped across sequential regions (Hd1-Hd4), with an increasing trend in diameter within the same prints; **e)** *In situ* over and under estimation frequency obtained by the overlay analysis of the *in situ* and *ex situ* masks; **f)** Systematic *in situ* over and under estimation with threshold of >80%.

Box plots of sequential diameters were generated for both vertical (Figure 4b) and horizontal (Figure 4d) orientations. All diameters were standardized using the dataset’s mean and standard deviation to identify deviations. A clear upward trend emerged within each print: early segments (e.g., Vd1) showed negative deviation, while later segments (e.g., Vd5) showed positive deviation. This was statistically significant for vertical filaments (p < 0.0001), though no statistical inference could be drawn for horizontal filaments despite a noticeable upward trend. The progressive diameter increase likely results from gradual pressure buildup in the extrusion system. This effect appeared consistent across both orientations, though shorter horizontal filaments, with higher velocity transitions, were less affected, likely due to non-stationary conditions.

Finally, the overlay analysis of the segmented regions - obtained by combining all non-spilled prints from the *in situ* monitoring system and comparing them against the corresponding *ex situ* masks - revealed recurring geometric deviations in specific areas of the constructs (Figure 4e). This overlay represents the spatial difference between the predicted and actual printed regions, effectively highlighting areas of under- and over- prediction by the system. The consistent patterns of deviation across multiple prints show the systematic print errors rather than random noise, underscoring the value of the *in situ* monitoring system for spatial error mapping and quality control (Figure 4f). Notably, these deviations were particularly pronounced in sections where the print head slowed down in the transitions between straight and curved regions, which likely caused greater disturbance in the support bath and amplified local optical distortions.

## Discussion

We present a complete *in situ* monitoring solution that combines a low-cost sensing architecture with a tailored data mining procedure. Embedded bioprinting presents unique challenges for *in situ* monitoring, as the overlying support bath can obscure the printed geometry, making it difficult to assess fidelity and resolve shape features accurately. We’ve addressed this through extensive prior testing, where we systematically evaluated multiple sensing modalities—thermal imaging, polar imaging, and visible imaging—and found that only visible imaging was effective in identifying anomalies and reconstructing local deviations from the nominal shape. This integrated approach enables the investigation of process parameter effects and stability across different locations within the bioprinted construct. Unlike methods proposed in literature, our system supports precise, real-time data collection during embedded bioprinting without the need for specialized or expensive equipment. The total cost of the imaging setup including the digital microscope, holder, and backlight was under$500, in contrast to systems such as OCT, which can cost upwards of$30,000.

On the data mining side, we also explored different segmentation strategies—including threshold-based methods, Otsu’s method, and a transformer-based image segmentation approach. Analyses showed that SAM-based segmentation consistently outperformed traditional thresholding, with higher correlation coefficients and reduced sensitivity to contrast and curvature. Both methods slightly overestimated *in situ* diameters, likely due to optical distortions from imaging through the support. Still, SAM’s strong agreement across conditions highlights its potential to improve measurement reproducibility. Moreover, SAM was adopted using its point-based prompt, which aligns well with the G-Code foundation of the bioprinting process, making it inherently compatible with any 3D-printable geometry and broadly adaptable across designs. The system enabled in-depth analysis of the printing process, identifying a critical threshold printing velocity for stable printing, while it was observed how diameter increases during print likely due to extrusion pressure buildup. Overlay analysis exposed recurring geometric deviations, confirming the system’s utility in mapping spatial errors.

While our system was optimized using a single bioink and support bath formulation—a bioink containing phenol red (a standard cell culture media additive) in a Pluronic-laponite matrix—it serves as a proof of concept for an *in situ* monitoring method that works with any bioink formulated in standard cell culture media. Given the diversity of bioinks and support materials in the field, our goal was to demonstrate effective monitoring under biocompatible conditions using phenol red, a contrast agent with an established history of cytocompatibility.

Beyond these findings, the modular architecture of the *in situ* monitoring system offers a versatile platform adaptable to various sensors, including different digital microscopes and advanced imaging cameras. This flexibility supports use across diverse bioprinting setups and materials, extending its utility beyond the current study. Design trade-offs such as balancing resolution, processing speed, and cost will be important as this system scales. Recognizing these trade-offs can help guide how similar sensing-AI approaches are adapted across biomanufacturing device applications.

Importantly, this system also supports two promising approaches to process optimization and control. First, in-situ process optimization can be achieved by using a small number of sacrificial prints, as demonstrated in this study, to empirically determine the optimal printing parameters, such as extrusion pressure and speed, for specific geometries or materials. Once calibrated, these parameters can be applied reliably in subsequent production runs. Second, the system provides a foundation for future studies in closed-loop process control. For example, printing may begin with a low extrusion pressure while real-time monitoring tracks local deviations from nominal filament dimensions (e.g., backside diameter). The system can then incrementally adjust the pressure until deviations fall below a predefined threshold. This approach assumes the availability of prior experimental mapping between pressure and printed diameter, and it opens the door to adaptive printing strategies, such as varying pressure across the construct to compensate for transient effects. Looking ahead, integrating closed-loop control capabilities supported by proof-of-concept demonstrations or simulations could advance this platform from a monitoring tool to a fully responsive control system.

These considerations strengthen the framing of our method as not only a monitoring tool but also as a robust foundation for intelligent process control in embedded bioprinting. By enabling real-time inspection, adaptive correction, and empirical parameter tuning, we anticipate that our approach can improve reproducibility, reduce material waste, and accelerate process optimization for real-world applications in tissue engineering.

## Materials and Methods

### Support Bath Preparation

The support bath was prepared according to our previous work ^30^. The final concentration of the support bath is 3% Laponite RDS (BYK Additives & Instrument), 10% Pluronic F127 (Sigma Aldrich), 0.5% calcium chloride (Sigma Aldrich), and 20 µl/ml of thrombin (Sigma Aldrich).

### Bioink Preparation

Sodium alginate (Sigma Aldrich) was mixed in Dulbecco’s Modified Eagle medium (DMEM, thermoFisher) thoroughly to achieve 1,5% (w/v) concentration. Fibrinogen from bovine plasma (Sigma Aldrich) was then added to the solution at a concentration of 6 mg/ml, and vortexed until a homogenous ink is achieved.

### 3D Bioprinting

A BIO X^™^ bioprinter (CELLINK®, Sweden) with a pneumatic extrusion printhead and a 0.33 mm needle-equipped cartridge was used. All printing consumables were sourced from CELLINK®.

### In Situ Imaging

All the *in situ* images of the printed geometry were captured using a Dino-Lite AM4113ZtL digital microscope. The custom holder was designed in Fusion 360 and 3D printed (Bambu Lab X1-Carbon, 0.4 mm layer height, 20% infill, PLA filament). A USB-powered LED backlight (MSAK827, DUNWELL TECH) was placed beneath the Petri dish during printing and imaging. Images (1280 × 974 px) were taken at 1/30 s exposure, resulting in 18 RGB *in situ* images.

### Ex Situ Imaging

A Nikon AXR laser scanning confocal microscope with a 4× objective was used (excitation: 488 nm; emission: 518–572 nm). For each construct, Z-stacks of 25 slices were acquired at 32 µm steps across the full sample thickness, with constant laser power and gain for consistency. With the Z-stacks, maximum intensity projections were then generated in ImageJ (Fiji). In total, 18 RGB images (5940 × 5940 px) were collected.

### Image Analysis

*In situ* RGB images were converted to grayscale, equalized, and segmented using a threshold of 235, with pixels with intensity ≥235 set as foreground. Masks were denoised via morphological closing (5×5 kernel) and removal of connected regions <500 pixels in area. *Ex situ* images followed a similar process, with added background removal (pixel-wise subtraction of a Gaussian-blurred 25×25 copy of the image) and re-equalization, followed by thresholding and denoising (removal of connected regions <3000 pixels in area, closing with 10×10 kernel). For SAM segmentation, the Vit-H model (sam_vit_h_4b8939.pth) was used via the official API in point-prompt mode: positive/negative points were placed, and the model returned binary masks for both *in situ* and *ex situ* images. *In situ* masks (1280×974) were registered to *ex situ* images (5940×5940) using annotated landmarks and a projective transformation.

### IMARIS Surface Reconstruction

For the IMARIS surface reconstruction, a high-magnification z-stack was acquired at the center of each geometry using the same confocal microscope as for *ex situ* imaging, with a 10x objective. The same excitation/emission settings were used as for the *ex situ* images. Each z-stack consisted of 134 images collected through the full sample thickness with a 6 µm step size. The stacks were then processed in IMARIS (version 10.2) using the surface reconstruction tool.

## Acknowledgements

The authors thank the Safety Health Environmental Discovery (SHED) Lab at MIT for providing equipment and workspace, as well as Dr. Sina Kheiri for helpful technical discussions related to this work. The authors acknowledge funding support from the Progetto Roberto Rocca Fellowship (awarded to G.Z.), the DoD Army Research Office Early Career Program and PECASE (W911NF-22-1-0126, awarded to R.R.) and the DoD DURIP Program (W911NF-24-1-0106, awarded to R.R.).

